## Supplemental Figures 1-6 for "TaRTLEt: Transcriptionally-active Riboswitch Tracer Leveraging Edge deTection"

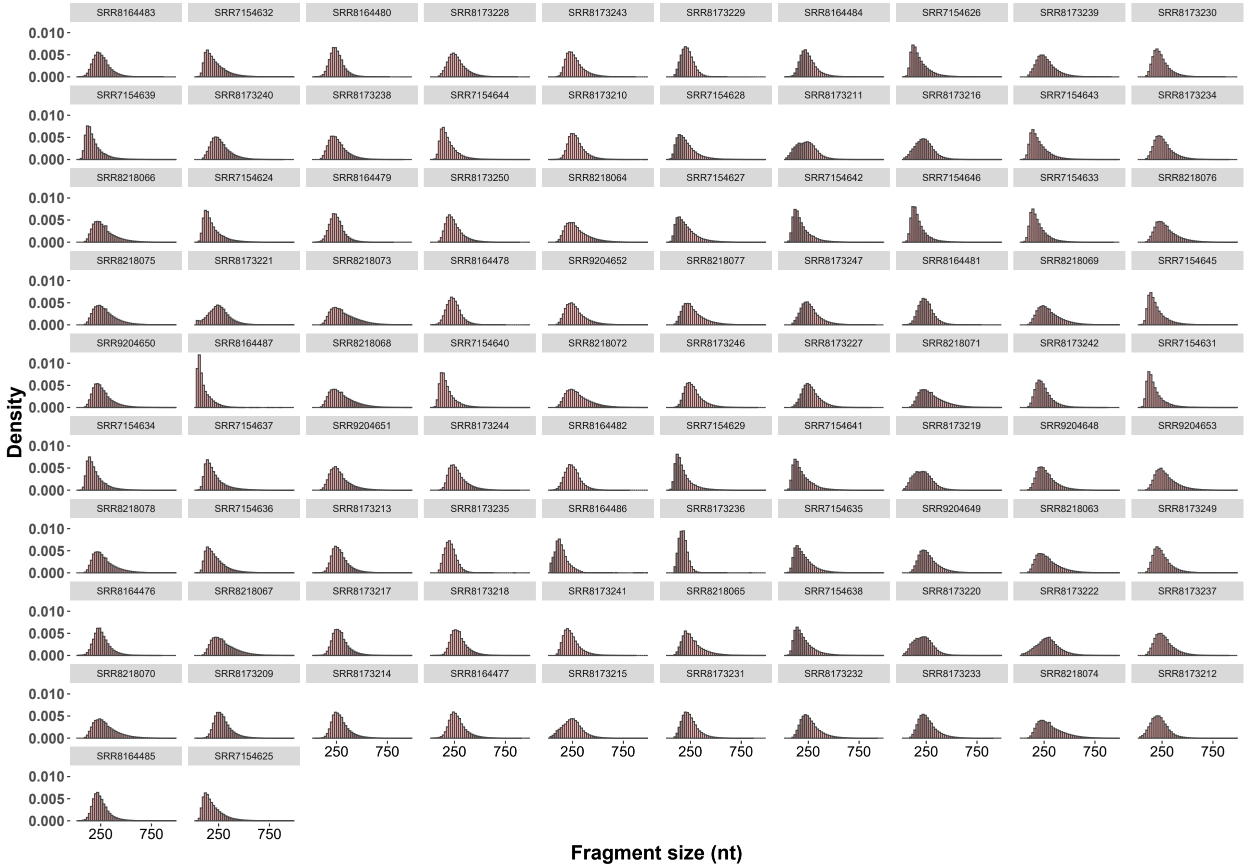


Figure S1: Read fragment length distributions for the 92 *E. coli* transcriptomes included in the `TaRTLEt` validation dataset.

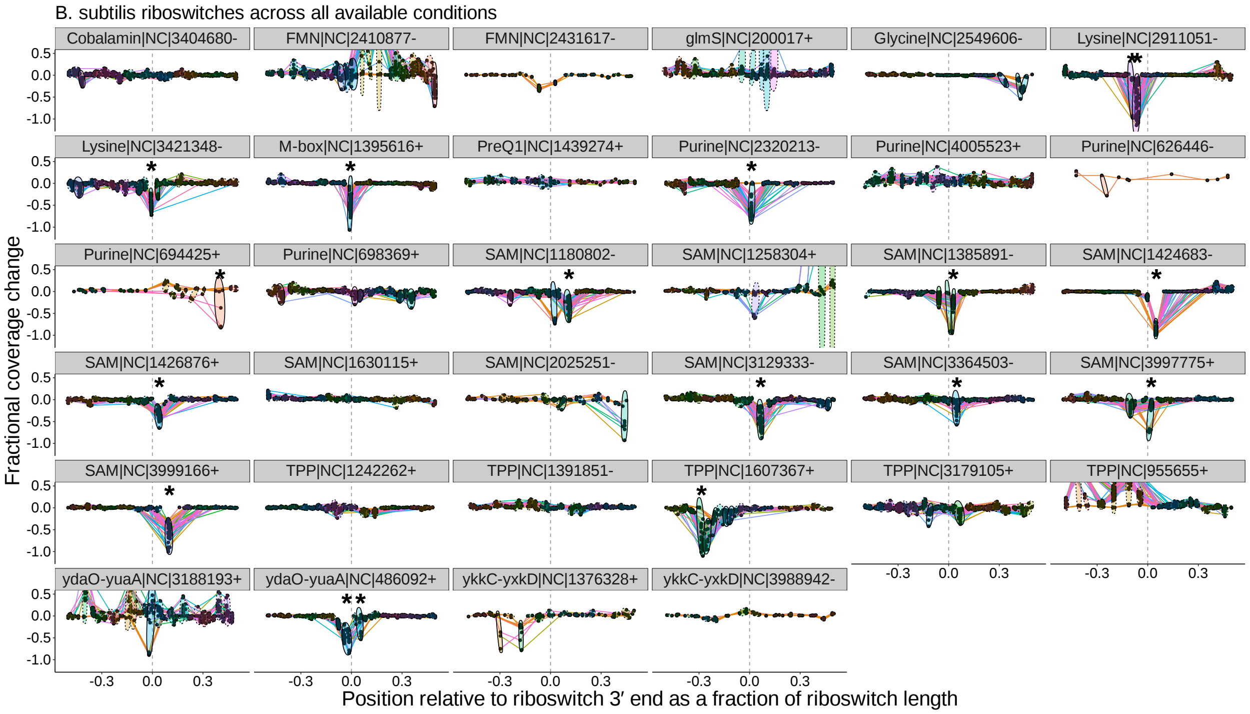


Figure S2: Peak plots for the 34 riboswitch loci identified by Infernal in the *B. subtilis* genome. Peak clusters are grouped by colored ellipses; clusters with at least one significant transcription-termination peak are bounded by solid ellipses, and those with no significant peaks are bounded by dashed ellipses. Asterisks mark peak clusters that (i) contain at least one significant transcription-termination peak, (ii) have significantly lower mean ($p < 0.05$, Mann-Whitney one-tailed U-test) than the set of all other peak clusters, and (iii) have significantly higher variance ($p < 0.05$, Levene's test) when compared to randomly sampled sets of the same number of peaks.

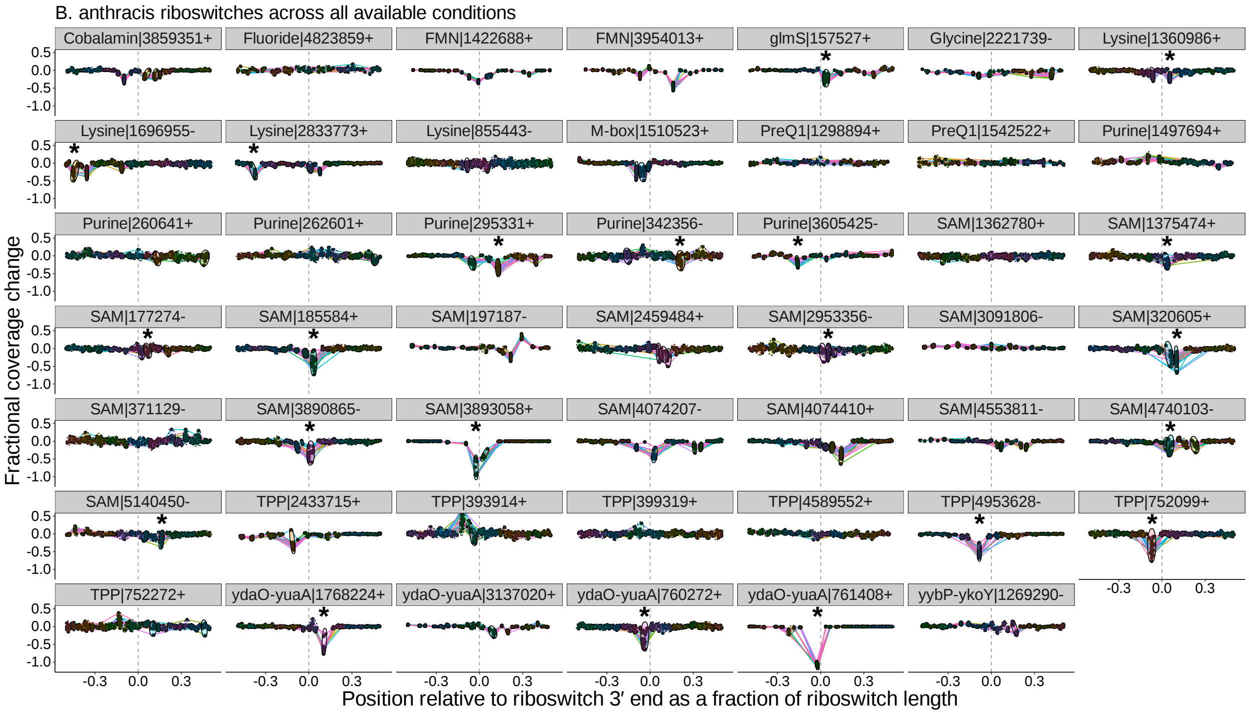


Figure S3: Peak plots for the 48 riboswitch loci identified by Infernal in the *B. anthracis* genome. Peak clusters are grouped by colored ellipses; clusters with at least one significant transcription-termination peak are bounded by solid ellipses, and those with no significant peaks are bounded by dashed ellipses. Asterisks mark peak clusters that (i) contain at least one significant transcription-termination peak, (ii) have significantly lower mean ($p < 0.05$, Mann-Whitney one-tailed U-test) than the set of all other peak clusters, and (iii) have significantly higher variance ($p < 0.05$, Levene's test) when compared to randomly sampled sets of the same number of peaks.

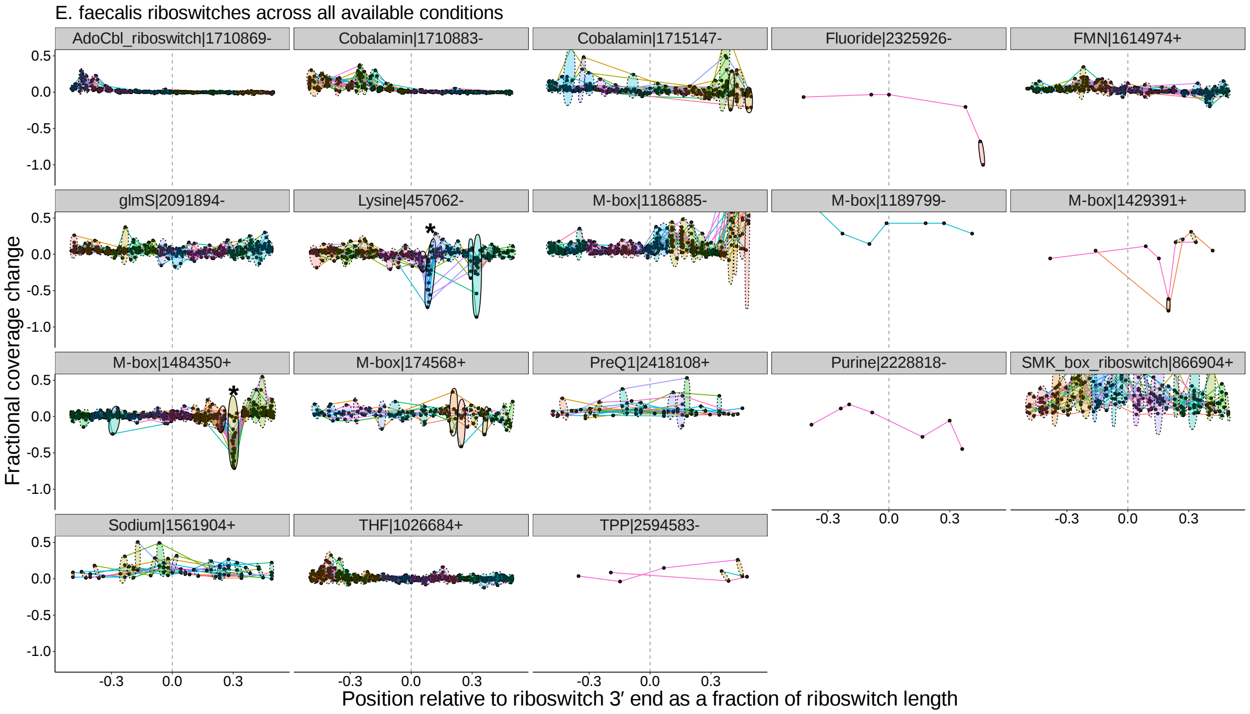


Figure S4: Peak plots for the 18 riboswitch loci identified by Infernal in the *E. faecalis* genome. Peak clusters are grouped by colored ellipses; clusters with at least one significant transcription-termination peak are bounded by solid ellipses, and those with no significant peaks are bounded by dashed ellipses. Asterisks mark peak clusters that (i) contain at least one significant transcription-termination peak, (ii) have significantly lower mean ($p < 0.05$, Mann-Whitney one-tailed U-test) than the set of all other peak clusters, and (iii) have significantly higher variance ($p < 0.05$, Levene's test) when compared to randomly sampled sets of the same number of peaks.

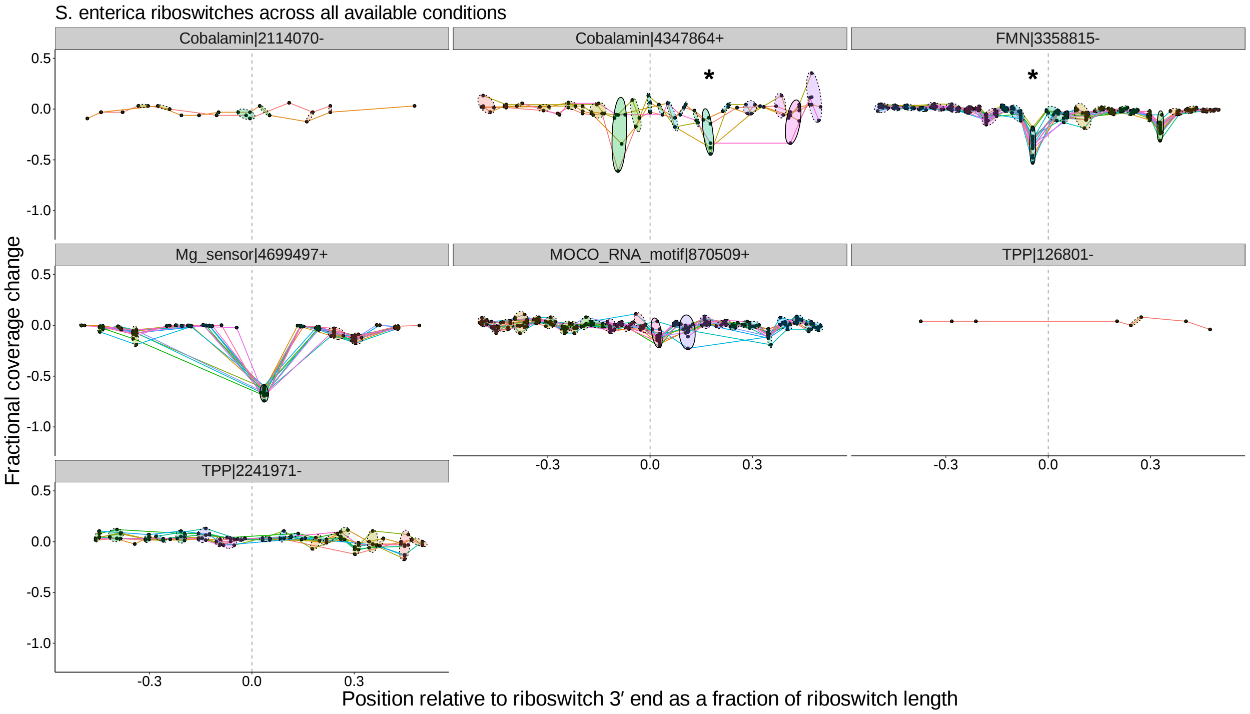


Figure S5: Peak plots for the seven riboswitch loci identified by Infernal in the *S. enterica* serovar Typhimurium genome. Peak clusters are grouped by colored ellipses; clusters with at least one significant transcription-termination peak are bounded by solid ellipses, and those with no significant peaks are bounded by dashed ellipses. Asterisks mark peak clusters that (i) contain at least one significant transcription-termination peak, (ii) have significantly lower mean ($p < 0.05$, Mann-Whitney one-tailed U-test) than the set of all other peak clusters, and (iii) have significantly higher variance ($p < 0.05$, Levene's test) when compared to randomly sampled sets of the same number of peaks.

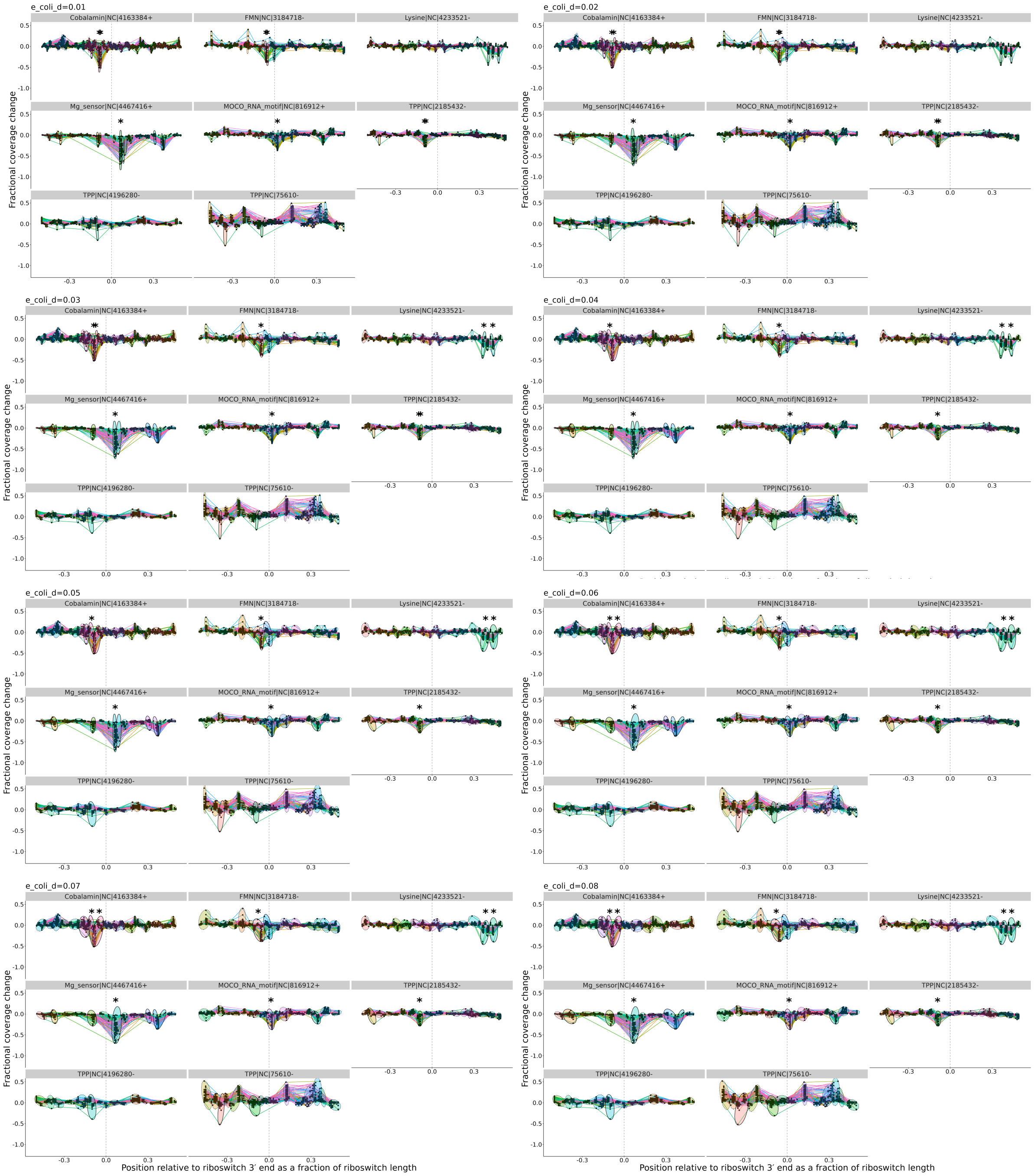


Figure S6: Sample peak plots guiding choice of clustering cophenetic distance threshold. Threshold values were scanned from 0.01 to 0.08 to examine clustering behavior and guide the choice of default. The resulting peak plots for *E. coli* are shown with the candidate threshold values increasing from left to right and top to bottom. A default value of 0.04 was chosen as well-behaved across the large majority of loci.
